# The role of the vestibular system in value attribution to positive and negative reinforcers

**DOI:** 10.1101/2020.05.17.100255

**Authors:** Elvio Blini, Caroline Tilikete, Leonardo Chelazzi, Alessandro Farnè, Fadila Hadj-Bouziane

**Affiliations:** Integrative Multisensory Perception Action & Cognition Team (ImpAct), INSERM U1028, CNRS UMR5292, Lyon Neuroscience Research Center (CRNL), Lyon, France; University of Lyon, Lyon, France; Department of General Psychology, University of Padova, Padova, Italy; Hospices Civils de Lyon, Neuro-Ophthalmology and Neurocognition, Hôpital Neurologique Pierre Wertheimer, Bron, France; Department of Neuroscience, Biomedicine and Movement Sciences, University of Verona, Verona, Italy; National Institute of Neuroscience – Verona Unit, Verona, Italy; Hospices Civils de Lyon, Neuro-Immersion Platform, Lyon, France

**Keywords:** Reward, Conflict, Monitoring, Monetary loss, Motivation, Vestibular Stimulation, Interoception

## Abstract

Somatic inputs originating from bioregulatory processes can guide cognition and behavior. One such bodily signal, mostly overlooked so far, is represented by visuo-vestibular coupling and its alteration, which in extreme cases may result in motion sickness. We argued that the inherently perturbed interoceptive state that follows can be a powerful determinant of human motivated behavior, resulting in a blunted response to appetitive stimuli and an exaggerated response to noxious ones. We sought to assess such differential impact of visuo-vestibular mismatches on value through a task involving conflict monitoring. We therefore administered to 42 healthy participants a modified version of the Flankers task, in which distractors (arrows, pointing in either a congruent or incongruent direction) signaled the availability of monetary incentives (gains, losses, or neutral trials). While performing the task, participants received either galvanic vestibular stimulation (GVS), or sham stimulation. We have found impaired behavioral performances when value, which was attached to task-irrelevant information, was at stake. Gains and losses, interestingly, dissociated, and only the latter caused enhanced interference costs in the task, suggesting that negative incentives may be more effective in capturing human attention than positive ones. Finally, we have found some weak evidence for GVS to further increase the processing of losses, as suggested by even larger interference costs in this condition. Results were, however, overall ambiguous, and suggest that much more research is needed to better understand the link between the vestibular system and motivation.

**Highlights:**

- Visuo-Vestibular mismatches may be important somatic markers affecting the evaluation of reinforcers;
- When attached to distractors, value information impairs behavioral performance for the task at hand;
- Trials in which potential losses were at stake were associated with larger interference costs arising from conflicting information between the target and the flankers;
- GVS (Right-Anodal) may further increase the interference caused by losses, but the evidence in this respect was ambiguous and inconclusive;

## 1. Introduction

### 1.1. Interoception and motivation

Human decision-making processes can hardly be understood in full without taking into account one individual’s physiological state and needs (Berridge & Robinson, 2003; Craig, 2002; Critchley et al., 2004; Damasio, 1996; Gray & Critchley, 2007; Namkung et al., 2017; Naqvi et al., 2007; Naqvi & Bechara, 2010; Paulus, 2007). “Somatic markers” originating from bioregulatory processes can bias the subjective desirability of stimuli, thus guiding cognition and behavior (Craig, 2002; Damasio, 1996; Mayer, 2011; Morton et al., 2006; Namkung et al., 2017; Paulus, 2007; Seth, 2013). For example, think how desirable would be one’s preferred food in normal conditions versus while experiencing mild nausea. A global interoceptive state is defined by the balance and integration of visceral, autonomic, somatosensory, motor and vestibular inputs, wherein the insula is known to represent a key structure in the integration of these signals (Craig, 2002; Namkung et al., 2017). The incentive value of any stimulus can be encoded in an abstract fashion, but also in relation to the expected effect on physiological homeostasis (Berridge & Robinson, 2003; Palminteri et al., 2012), before being exploited for guiding human choices via connections with other structures such as limbic areas and the dorsal striatum (Berridge & Robinson, 2003; Namkung et al., 2017; Palminteri et al., 2012). This double coding of one stimulus (i.e. abstract vs. homeostasis-related) is therefore capable to explain why the same stimulus can assume different motivational values under different interoceptive states.

### 1.2. The vestibular system: a sensory entry to interoception and motivation

One important, yet overlooked, bodily signal is the uncoupling of normally tightly linked inputs, i.e. visual and vestibular information, which in the most extreme case results in motion sickness and nausea (Kohl, 1983; Lackner, 2014; Treisman, 1977). Motion sickness is a complex syndrome which manifests itself not only with nausea and vomiting but also with a variable degree of pallor, cold sweating, drowsiness, and dizziness. Though the physiological origins of motion sickness are not yet fully understood, the most widely accepted theory posits that sensory conflicts (i.e. a discrepancy between the expected and afferent signals from the body) may provoke the extreme reaction of neural mechanisms in the brainstem and several cortical regions (Lackner, 2014), including responses of anxiety and aversive conditioning (Balaban, 2002). A partly alternative, evolutionary account posits that motion sickness may essentially be related to poisoning (Treisman, 1977). Under this evolutionary framework, visuo-vestibular mismatches are reminiscent of the subtle but informative warning signals which follow the ingestion of noxious neurotoxins (Treisman, 1977), and thus would prompt defensive reactions such as emesis. Furthermore, such mismatches would participate to the process of aversive conditioning in the sense that they would provide to an unspecialized feeder elements for avoiding potentially noxious stimuli or environments in the future (Treisman, 1977). At any rate, the experience of visuo-vestibular mismatches may be regarded as a powerful perturbation of the human interoceptive state.

Vestibular stimulation techniques – which alter the activity of an extended parieto-insular network (Lopez et al., 2012; zu Eulenburg et al., 2012) – have been found to modulate affective control, mood, purchase decision-making (in terms of desirability of products) in healthy subjects (Mast et al., 2014; Preuss et al., 2017; Preuss, Hasler, et al., 2014; Preuss, Mast, et al., 2014), and to alleviate manic symptoms in patients (Carmona et al., 2009; Dodson, 2004; Levine et al., 2012). For example, Preuss, Hasler, et al. (2014) reported that caloric vestibular stimulation can modulate the performance in a Go/No-go task exploiting emotional images as target stimuli. In their study, affective control (i.e. the proportion of *hits* vs. *false alarms*, and thus motor inhibition) for positive images was modulated by the vestibular stimulation, and could either be reduced or increased as a function of the stimulated ear. Moreover, one recent study found that vestibular stimulation through galvanic current (Galvanic Vestibular Stimulation, GVS) decreases sensitivity to monetary rewards (Blini, Tilikete, Farnè, & Hadj-Bouziane, 2018a). Therefore, stimulating the vestibular sense may be a key and convenient mean to perturb interoceptive states or otherwise compensate for a system that does not link efficiently visceral states to optimal decision-making strategies, as for instance in addicted individuals.

### 1.3. An eye on behavioral addictions

In addiction disorders, cues associated with one’s behavioral addiction exert a strong attentional capture (Field, Munafò, & Franken, 2009). The magnitude of attentional bias has been found to predict relapse from treatment (Field & Cox, 2008; Garland, Franken, & Howard, 2012; Marissen et al., 2006), and has been causally related to craving (Field & Eastwood, 2005). Such cues activate automatic representations of value which can interfere with the task at hand, when they should rather be inhibited because distracting (Carpenter et al., 2006; Cox et al., 2006; Gross et al., 1993; Nijs et al., 2010; Sokhadze et al., 2008). For example, in the manifold variants of the Addiction-Stroop test (Cox et al., 2006), participants with behavioral addictions are typically found to be slower than controls in reading aloud the color of words related to the substance of abuse. Thus, interference costs can provide valuable clinical information, on one hand, and reducing them could represent a major therapeutic objective, on the other hand. This appears particularly appropriate as prominent models of behavioral addictions emphasize – among other societal, neurobiological, and cognitive aspects – the role of both “driving” (enhanced sensitivity to reward) and control (reduced inhibitory control) aspects, and their interaction (Baler & Volkow, 2006; Della Libera et al., 2019; Goldstein & Volkow, 2011).

Emerging evidence also points toward an important role of defective interoceptive processing in craving and in the maintenance of behavioral addictions (Gray & Critchley, 2007; Paulus et al., 2013; Verdejo-Garcia et al., 2012). New avenues for the clinical management of these patients include treatment options based on interoceptive processing (Paulus, 2007; Paulus et al., 2013), their effectiveness being explained in light of a possible modulation of insular activity. Indeed, inactivation of the insula in amphetamine-experienced rats may prevent their drug-seeking behavior and blunt the malaise associated with craving (Contreras et al., 2007); in humans, damage to or functional disruption of the insula may alleviate addiction to nicotine (Dinur-Klein et al., 2014; Naqvi et al., 2007). Thus, manipulating this circuitry represents an appealing way to tackle the addiction loop in the brain (Dinur-Klein et al., 2014; Droutman et al., 2015; Naqvi et al., 2007; Naqvi & Bechara, 2010; Paulus, 2007). However, the effects of vestibular stimulation – which also taps onto an extended parieto-insular network (Lopez et al., 2012; zu Eulenburg et al., 2012) – have been seldom studied.

### 1.4. Assessing the role of the vestibular system in value attribution

Detrimental effects on performance and control induced by value-associations have also been described in healthy individuals, and characterized as being a function of the distractors’ salience and the degree of automaticity of their processing (Bourgeois et al., 2016; Chelazzi et al., 2013; Failing & Theeuwes, 2018; Krebs et al., 2010). This effect is important because it provides a working model to study a key feature of addiction disorders in a laboratory setting. In such experimental conditions, value-associations are typically established using monetary rewards. With this study, we therefore sought to test, in healthy subjects, whether a vestibular/interoceptive perturbation is capable to decrease the unduly interference induced by intrinsically salient features (Krebs et al., 2010). Second, we sought to test whether such artificially biased interoceptive state differentially affects rewards and punishments. Whether negative reinforcers are devaluated following visuo-vestibular mismatches, as are positive ones, is currently unknown. One study found that inactivation of the posterior insula in rats may result in disrupted acquisition of both conditioned place preference and place avoidance (Li et al., 2013). Indeed, a perturbed system that strives for homeostasis may bring about a general blunted response to external stimuli, as physiologic balance would become the main priority; this predicts decreased sensitivity to punishments (Scenario 1 in **Figure 2C**). However, the same study also found evidence for a neural dissociation: lesions to the anterior insula selectively disrupted conditioned place avoidance, but not conditioned preference (Droutman et al., 2015; Li et al., 2013). One possibility is thus to observe a similar behavioral specificity. For example, in the study of Preuss, Hasler, et al. (2014), affective control was modulated for positive, but not negative images. Should the vestibular perturbation be specialized for appetitive stimuli, negative reinforcers may be unaffected (Scenario 2 in **Figure 2C**). However, differently from positive reinforcers, punishments have a threatening nature and negative valence, which appear to match the nature of the altered interoceptive processing and may resonate with it. As visuo-vestibular mismatches may be exploited as (further) warning signal (e.g. signaling the contact with neurotoxins, Treisman, 1977), negative reinforcers may become more salient and, ultimately, more effective in capturing attention. We therefore put forward the possibility that the effects of negative reinforcers may be enhanced by a vestibular stimulation (Scenario 3 in **Figure 2C**, and possible neural substrates in paragraph 1.5). In the context of addiction disorders, this would be equally desirable to a decreased sensitivity to positive rewards to the extent that, in the context of decision making, short and long term negative consequences are weighted comparatively more than short term positive effects, i.e. “myopia” for the future is counteracted (Bechara, Dolan, & Hindes, 2002).

We administered GVS to healthy participants engaged in a task evoking conflict and the need to inhibit irrelevant information (an ad-hoc adaptation of Eriksen & Eriksen, 1974; Eriksen, 1995). Participants received, in three different days, one sham stimulation and two active GVS stimulations, with opposite hemispheric lateralization (Blini, Tilikete, et al., 2018a). We used a modified version of the Flankers Task (FT), whereby five arrows were presented on screen: the task consisted in indicating in which direction the central arrow was pointing. The four flanking arrows could either point toward the same (congruent) or the opposite direction (incongruent), thereby creating interference costs in the incongruent condition. Depending on their color, flankers could also signal the possibility of receiving monetary incentives or losses, as a function of the subjects’ performance. Interference costs were expected to increase when – as compared to a Neutral (N) condition without reward – Potential Gains (PG) or Potential Losses (PL) were at play (Krebs et al., 2010), reflecting increased attentional processing. We had two predictions: 1) interference costs would be maximal, in PG conditions, for the sham stimulation, but reduced by GVS, especially with a Right-Anodal montage (which was associated with the greatest reduction in sensitivity to monetary rewards in a previous study, Blini, Tilikete, et al., 2018a); 2) interference costs for PL, on the contrary, would be enhanced with either GVS condition with respect to the sham stimulation (Scenario 3 in **Figure 2C**).

### 1.5. Potential neural mechanisms

In **Figure 1,** a schematic depiction of a neural mechanism possibly involved in the link between vestibular perturbations and motivation is presented, largely inspired by the Embodied Predictive Interoception Coding (EPIC) model of Barrett & Simmons (2015). The EPIC model stems from recent active inference accounts (e.g. Adams, Shipp, & Friston, 2013), holding that perception is a process of statistical inference and prediction about the causes of sensations (also see Seth, 2013). Under this framework, the (external) context at hand would be first evaluated through a prior model, based on past experience, about the most likely output under the circumstances. This comparison allows the brain to issue timely predictions about the most appropriate bodily response, the required allostatic adjustments, and the interoceptive sensations that are expected to arise. These predictions are thus the basis for the perception of and the action toward the external context, which can modify the context in turn (e.g. by modulating attentional resources, or by prompting approach-withdraw behaviors). A widespread network of visceromotor cortices – e.g. posterior orbitofrontal cortex, posterior ventral medial prefrontal cortex, anterior insula, and anterior cingulate cortex – has been putatively ascribed to this function: these areas (agranular cortices) present a laminar architecture characterized by a large number of pyramidal neurons in the deepest layers in face of fewer projection neurons in the upper layers, a feature that suggests a role in sending sensory predictions to granular cortices, which are better equipped for computing prediction errors. Indeed, the “primary interoceptive cortex”, i.e. the granular posterior insula, appears as one key region in which these predictions are compared to the actual incoming sensations. If predictions are roughly accurate, the difference will be null or negligible; if large discrepancies between predictions and actual sensory signals are, instead, detected, one prediction error signal will back-propagate to visceromotor cortices and modulate their activity. As a result, the prediction error may adjust predictions and the prior model that generated them directly, in both cases affecting the actions the body will undertake and the evaluation of the context.

**Figure 1:**
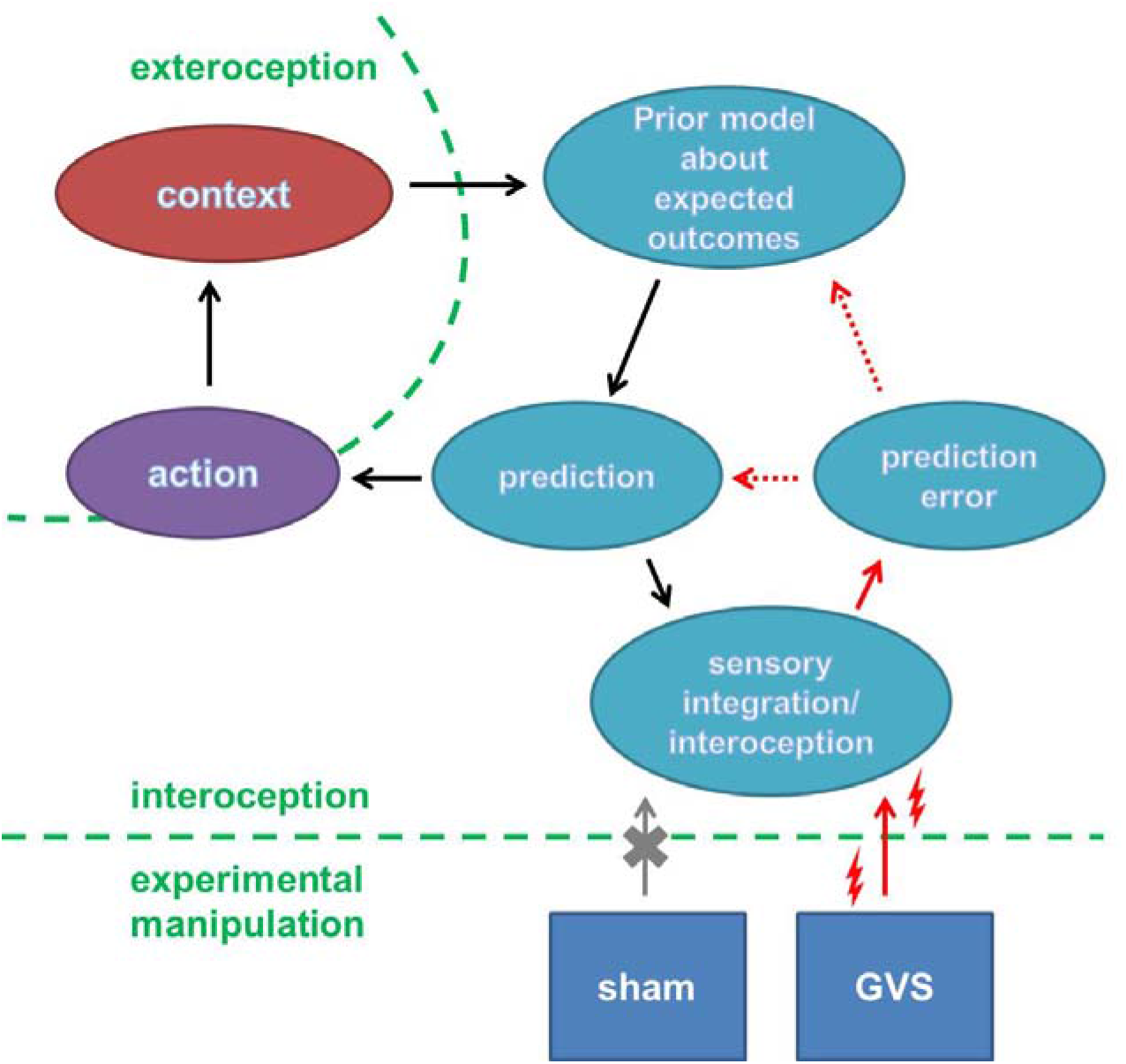
schematic depiction of a mechanistic model possibly involved in the link between vestibular perturbations and the external context (e.g. monetary gain vs. losses), largely inspired by the Embodied Predictive Interoception Coding (EPIC) model of Barrett & Simmons (2015). The context at hand would be first evaluated through a prior model, based on past experience, about the most likely output under the circumstances. This comparison allows the brain to issue timely predictions about the most appropriate bodily response, and the interoceptive sensations that are expected to arise. These predictions are thus the basis for the perception of and the action toward the external context, which can modify the context in turn. The prediction would be then compared to the actual incoming sensations in the primary interoceptive cortices. If large discrepancies between predictions and incoming sensory signals arise, one prediction error signal will back-propagate (dashed arrows) to adjust predictions and the prior model that generated them. This in turn would affect the actions the body will undertake and the subsequent evaluation of the context. By biasing the interoceptive system toward a more negative state, GVS, as opposed to sham, may increase prediction errors. While a positive context (e.g. monetary gain) may be devaluated, a negative one (e.g. monetary loss) could be exaggerated instead.

Under this framework, we speculated that, while the sham stimulation would not (sensibly) interfere with the process of sensory integration, vestibular stimulation would bias the incoming sensory signals toward a negative and unpleasant bodily state, in light of visuo-vestibular conflict. As a result, prediction errors would be larger than what expected on the basis of previous experience, and will guide the subsequent predictions toward a more negative valence attribution. The responses of the brain may therefore adapt differently to the context at hand. Although the present study did not envisage neural measures able to test this mechanistic model specifically, we predicted that a positive context, such that arising from monetary rewards, would be resized and devaluated (Blini, Tilikete, et al., 2018a); a negative context, arising when potential losses are at stake, would be magnified instead. This further motivated our prediction (**Scenario 3** in **Figure 2C**) of enhanced interference costs for PL in face of reduced interference costs for PG with concurrent GVS.

**Figure 2:**
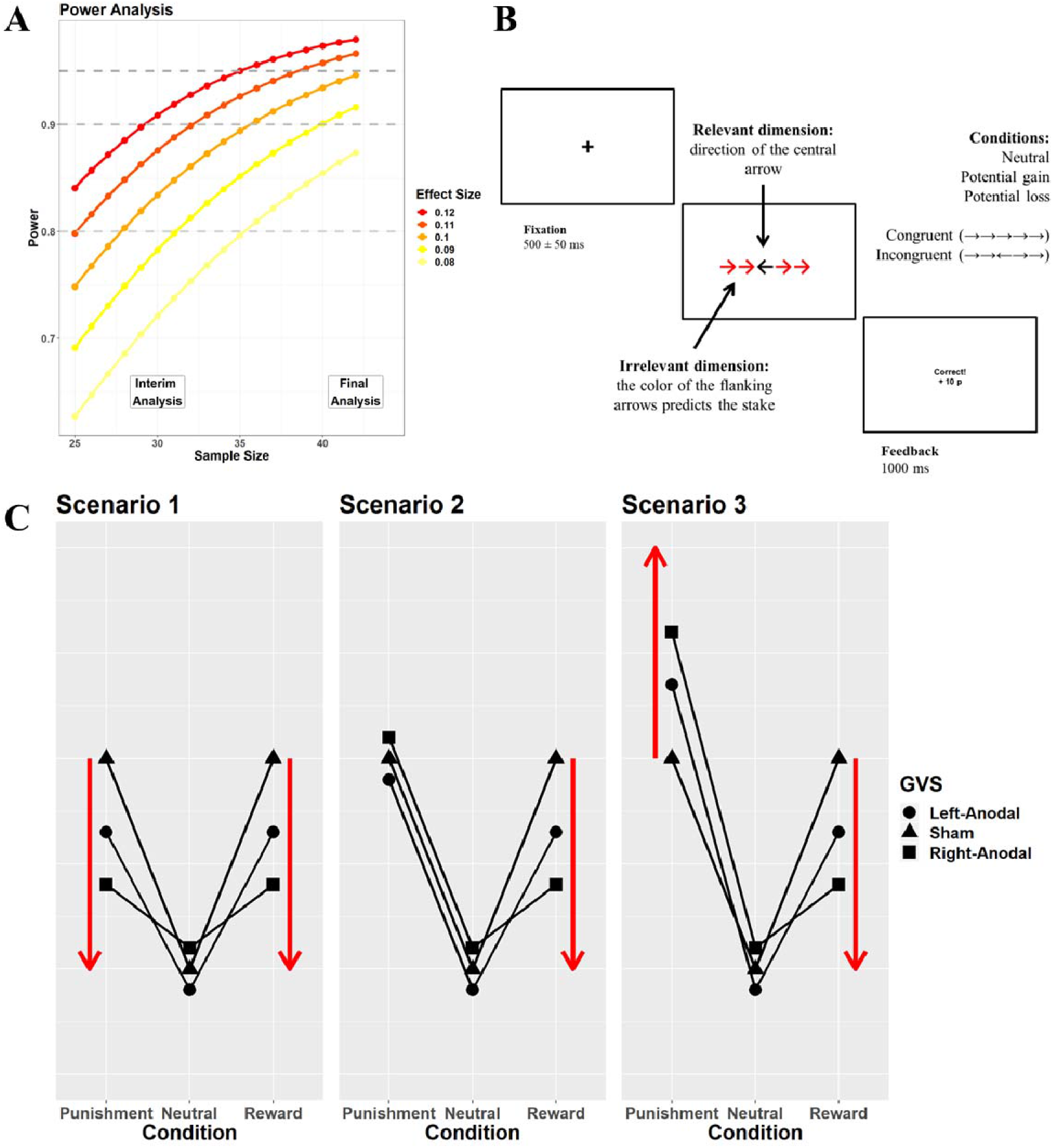
**A)** The nominal power for our design is depicted as a function of a range of a priori sample and effect sizes (partial eta squared). We planned to enroll a maximum of 42 participants, which allowed us to have high power for effect sizes in the small range (distributed around that reported in Blini et al., 2018a). However, interim analyses were planned at N= 30, and thus corrections for sequential analyses were applied. **B)** Graphical representation and time course of the modified Flankers Task (FT). Each trial started with a fixation cross appearing at the center of the screen. Then, a set of five arrows was presented on screen. The task consisted in indicating the direction of the central arrow (left, right). The four flanking arrows could point towards the same (congruent condition) or opposite direction (incongruent condition), the latter creating conflict and interference costs. Furthermore, the color of flanking arrows (the irrelevant dimension) was associated with one of three Conditions: Neutral (N, blue), in which no points were at stake; Potential Gain (PG, green), in which ten points (about 12 eurocents) were at stake; Potential Loss (PL, red), in which a 10 points loss was possible. A visual feedback was then presented on screen informing the participants about the outcome. The overall goal of this task was to highlight the attentional capture effects of value-associated features that, though irrelevant, were expected to interact with congruency and enhance interference costs. **C)** A priori possible scenarios for the effects of GVS on interference costs caused by Potential Gains (PG) or Potential Losses (PL). We predicted reduced sensitivity to rewards during GVS, which may reduce interference costs in the PG condition, especially with a Right-Anodal montage (Blini et al., 2018a). Different scenarios were then hypothesized, depending on the effects of GVS in the PL condition. The first possibility (Scenario 1) was that GVS could reduce sensitivity to punishments as well, and thus reduce interference costs in the PL condition. While the body is experiencing visuo-vestibular mismatches, response to external stimuli may be globally reduced, as physiologic balance would become the main priority and would subtract attentional resources to the external context. Scenario 2 assumed a weak dissociation between the treatment of PG and PL, as to accommodate the manifolds studies reporting partially distinct neural substrates and behavioral effects of reward and punishments (e.g. Cubillo, Makwana, & Hare, 2019). However, we favored a strong dissociation (Scenario 3) according to which interference costs would be magnified by GVS for PL. We argued that the negative valence of punishments may match the nature of the altered interoceptive processing and, differently from rewards, gain in salience (also see **Figure 1**, for a mechanistic model).

## 2. Methods

The registered protocol (https://osf.io/2htd3) and all supporting materials (**Supplementary Materials:** https://osf.io/b9ezq/) are available on the Open Science Framework website.

### 2.1 Participants

We computed the nominal power for a 2×3×3 general linear model design (F test, fully within subjects, alpha= 0.05) for a range of sample and effect sizes (**Figure 2A**, details and script in the **Supplementary Materials**). We decided to enroll a maximum of 42 participants. This sample size allowed us to have an adequate nominal power (94.6%) for effect sizes that we judged to be small, distributed around that reported in our previous unbiased study about GVS and reward processing (η_p_^2^= 0.1, Blini, Tilikete, et al., 2018a). We planned, however, an intermediate analysis at N= 30 (power: 83.4%). To avoid the spread of false positives, our threshold for significance was modified in order to preserve a cumulative false discovery rate of 0.05 (Lakens, 2014). Pocock’s correction for two sequential analyses suggested adjusting the alpha level to 0.034 (Pocock, 1977). Furthermore, p-values above 0.97 would have been indicative of futility of further data acquisition. Data collection would have been interrupted at N= 30 if one boundary was crossed. However, interim analyses provided, for our main tests, results that were only significant at the uncorrected threshold, and therefore we proceeded with recruiting 42 participants.

We recruited 42 young healthy participants (18-29 years, 18 males and 24 females, mean age: 22.3 years, SD: 2.8 years), which provided written informed consent. Inclusions were made on the basis of a medical examination aimed at checking inclusion and exclusion criteria. Inclusion criteria were: right-handedness; normal or corrected-to-normal vision. Exclusion criteria were: history of neurologic (e.g. epilepsy up to first degree of familiarity, migraines), psychiatric, cardiac, or otologic disorders (e.g. recurrent otitis media, perforation of the tympanic membrane); presence of metallic implants or splinters in the body; severe sleep deprivation, consumption of psychotropic drugs or alcohol during the previous 24 hours; participation to other brain stimulation experiments during the previous week. The study has been approved by the relevant French Institution (Comité de Protection des Personnes, CPP, 2015-A00623-46).

### 2.2 Galvanic Vestibular Stimulation (GVS)

GVS protocol mimics that described in Blini et al. (2018a). GVS was delivered *via* a commercial, CE approved, stimulator (BrainStim, EMS, Bologna). The application of small current intensities over the mastoid bones is associated with the illusion of head and body movements towards the side of anodal stimulation, but induces very few adverse effects with stimulation up to 1.5 mA in both healthy and brain-damaged patients (Utz et al., 2011). Electric current of 1 mA was administered continuously through spongy electrodes (14 cm^2 area) soaked with saline water and fixed in place with adhesive tape and a rubber band. Stimulation was delivered only after an initial impedance check, to minimize potentially painful sensations. Three configurations were adopted. Left-and Right-Anodal were active GVS conditions, inducing different (polarity dependent) effects. Electrodes were placed over the mastoid processes symmetrically (that is, in the Left-Anodal montage the cathode was placed over the right mastoid bone, and *vice versa* for the Right-Anodal montage). Left-Anodal stimulation activates mainly right hemisphere structures (Lopez et al., 2012; zu Eulenburg et al., 2012), whereas Right-Anodal activates comparatively more left hemisphere structures. A sham condition was also included, with electrodes placed symmetrically about 5 cm below the mastoids, above the neck, and distant from the trapezoidal muscles yielding proprioceptive signals (Lenggenhager et al., 2007). The sham condition was included to control for unspecific factors of electrical stimulation (e.g., arousal, discomfort). The anode was placed in this case on the left side (Ferrè et al., 2013). Participants performed the behavioral tasks three times, on three different days, under each GVS condition (Left-Anodal, Right-Anodal and sham). The order of GVS type administration was counterbalanced across subjects. Within each session, active stimulations were delivered for a maximum of 30 minutes in order to minimize side effects.

### 2.3 Apparatus and Behavioral Tasks

Participants were tested in a dimly lit, quiet room. Their head was restrained by a chinrest, facing a 17 inches large screen at a distance of approximately 57 cm. The open-source software OpenSesame (Mathôt et al., 2011) was used to display experimental stimuli on the screen and record the subjects’ response. Participants provided responses by means of keyboard presses (on a standard QWERTY keyboard) using the index and middle fingers of their dominant hand. During the stimulation, they performed two tasks. One control task **(Subjective Visual Vertical, SVV)** was administered during the first 5 minutes of the stimulation. SVV required rotating visual segments until they appeared to be in a vertical position. It was meant to provide independent evidence for a successful vestibular stimulation, given that displacements occur towards the site of anodal GVS stimulation (Mars et al., 2001; Saj et al., 2006). Then, a modified version of the **Flankers Task (FT)** (Eriksen & Eriksen, 1974; Eriksen, 1995) probed interference costs arising from conflict and their modulation by potential rewards or losses. Practice trials for both tasks were administered at the beginning of each session, before the onset of the stimulation, and were not considered in the analyses. One brief evaluation of subjective feelings and sensations experienced during GVS was administered soon after the stimulation end (see **Supplementary Materials**). This was meant to monitor participants’ distress and task compliance across the different days of the experiment and GVS protocols.

#### 2.3.1 Subjective Visual Vertical (SVV)

The SVV task is depicted in **Figure 3**. One segment (19 cm, 2 mm wide; white-colored, over a black background) was presented at the center of the screen. Its starting orientation varied randomly between 1 and 20 degrees from the geometric (objective) vertical, in both clockwise and counter-clockwise directions (counterbalanced). Participants were asked to align the segment along the vertical plane by manually rotating it clockwise or counter clockwise using the keyboard. Segments’ orientation changed in steps of 0.1° (between −1.6 and 1.6 degrees from the vertical) up to steps of 0.7° for more extreme responses (>19°). Participants were informed that a counting strategy was counter-productive, and they were asked to stress accuracy over speed.

**Figure 3:**
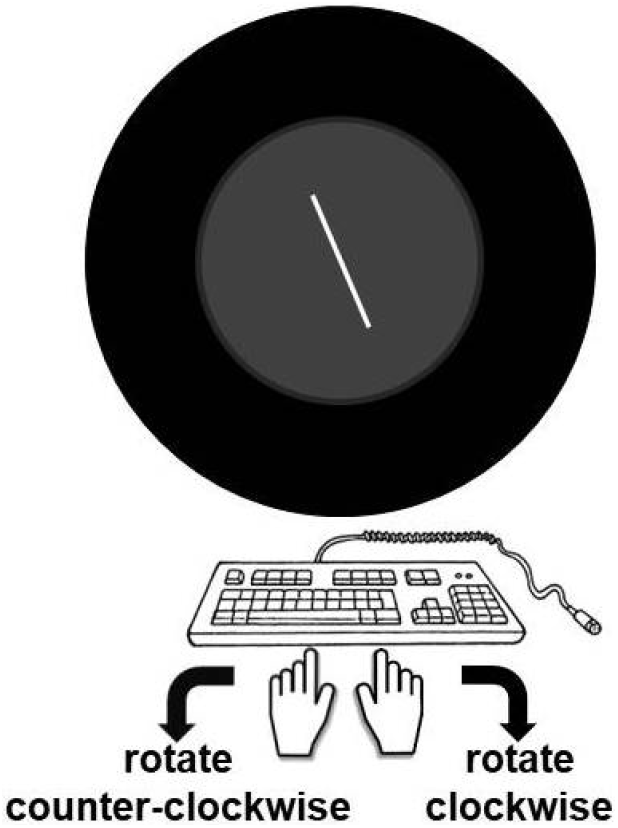
Graphical depiction of the SVV task. Participants were asked to align a visual segment to the perceived vertical orientation by means of manual responses. The task was performed in darkness and with a circular black panel covering the screen edges, as to minimize external visual anchors. Because GVS affects the inputs from the otoliths, which appear to play a key role in this task, biases were expected to occur in the direction of the anodal site of stimulation (i.e. clockwise biases for Right-Anodal, and counter-clockwise for Left-Anodal). The task was thus administered as a control aimed at confirming that the vestibular system was, indeed, perturbed.

The task was performed in darkness and a circular black panel covered the borders of the screen, to minimize the use of external anchoring points. A total of 24 trials was administered. As dependent variable, the orientation of the SVV (in degrees) was stored. Positive values reflect a shift occurring clockwise (i.e. towards the right ear), while negative values reflect a counter-clockwise bias. The script for running the task is available in the **Supplementary Materials**.

#### 2.3.2 Flanker Task (FT)

A schematic depiction of the task is illustrated in **Figure 2B**, and the script for running it is available in the **Supplementary Materials**. Each trial started with a fixation cross (1.2×1.2°) appearing at the center of the screen for 500 ms (±50 ms of uniformly distributed jitter). Then, a set of five arrows was presented on screen (about 0.5° of width each) until the subjects’ response. The task consisted in indicating the direction of the central arrow (left, right, equiprobable), white colored over dark background. The four flanking arrows could point toward the same (congruent condition) or opposite direction (incongruent condition), the latter creating conflict and interference costs. Furthermore, participants were explicitly instructed that the color of flanking arrows (the irrelevant dimension: red, blue, or green) was associated with one of three Conditions: Neutral (N), in which no points were at stake; Potential Gain (PG), in which ten points (about 12 eurocents) were at stake; Potential Loss (PL), in which a 10-point loss was possible. The color-value associations (i.e. red for PL, blue for N, and green for PG) were kept constant across subjects and sessions; note that color-value associations were explicit, i.e. not established via statistical learning paradigms. Finally, following the response, a visual feedback was presented for 1000 ms. The feedback consisted of two lines of text: the first indicated whether the response was correct, incorrect, or too slow; the second one indicated the number of points gained or lost for the last response. Participants earned the points at stake in PG, or avoided losing them in PL, only if they could provide a correct response faster than 75% of their overall responses. Participants were informed about this adaptive speed threshold. We reasoned that a challenging speed threshold would maximize the chances of observing reward-related behavioral signatures by enhancing the salience of PG and PL conditions, and the urgency of providing a response in such cases. As the effect of reward can be framed as an increased vigor deployed in conditions requiring extra effort, little or negligible effects could be in principle expected with a long response time-window, in which performance boosts are not crucial in order to obtain the incentives at stake. On the contrary, a more challenging task could help avoid ceiling effects for variables such as accuracy, by favoring impulsivity and incorrect responses, and eventually magnify behavioral effects such as enhanced congruency costs in PG and PL conditions. The presence and magnitude of such effects would also help interpret possible modulations by GVS. Differently from previous approaches (e.g. Blini et al., 2018a), this threshold was not fixed, but adaptive, and it was meant to provide similar levels of task difficulty across all subjects (e.g. Shen & Chun, 2011). Participants received a monetary compensation of 75 euros for their participation in the three sessions. They were also able to receive an additional amount up to 25 euros/session according to their performance in the FT task, proportionally to the amount of points gained.

The FT consisted of 648 trials overall, with two shorts breaks, for a total duration that did not exceed 25 min. The overall design counted 108 observations per cell for the two main manipulated factors (Congruency and Condition, 2×3). It was repeated once for each one of the three GVS conditions (1944 trials overall).

### 2.4 Analyses

Data, excluding practice trials, were analyzed with the open-source software R (The R Core Team, 2018). The following packages greatly eased our work: afex (Singmann et al., 2019); dplyr (Wickham, François, Henry, & Müller, 2019); emmeans (Lenth, 2019); ggplot2 (Wickham, 2016); lme4 (Bates, Mächler, Bolker, & Walker, 2014). For exploratory analyses, we recovered the parameters of a drift diffusion model (Ratcliff, 1978) using the fast-dm-30.2 software (Andreas Voss & Voss, 2007).

We decided to analyze accuracy only if: the group mean would score below 90% (and below 95% in congruent trials); if fewer than half of the subjects would present more than 90% success rate (95% in congruent trials). This was to avoid, in the FT task, ceiling effects. The group mean was eventually 84.8% (91.2% for congruent trials). 13 subjects (31%) presented more than 90% success rate in the FT task; 17 subjects (40.5%) presented more than 95% correct responses in congruent trials. Thus, we proceeded in assessing accuracy as dependent variable.

We assessed response times only for correct responses. Response times slower than +2.5 and faster than −2.5 standard deviations from the subjects’ mean – separately for the levels of GVS, Congruency and Condition – were discarded (0.02% overall). All participants presented more than 70% of correct responses on average. This threshold, though arbitrary, was established a priori in an attempt to operationalize what should have been regarded as “poor performance”, suggestive of lack of compliance or engagement; any subject with less than 70% accuracy would have been excluded from analyses. Likewise, all participants could attend all the three experimental sessions, and could receive GVS without inconveniences (e.g. excessively high impedance, excessive discomfort). No replacement was thus necessary. Data were analyzed through *mixed-effects multiple regression models* (Baayen et al., 2008) using the lme4 package for R (Bates, Maechler, Bolker, & Walker, 2014). Models had a logistic link-function, appropriate for binary variables, when assessing accuracy. Prior to fixed-effect testing, the most appropriate and parsimonious (Bates, Kliegl, Vasishth, & Baayen, 2015) matrix of random effects was chosen via an objective pipeline exposed at length elsewhere (Blini, Desoche, et al., 2018b; Blini, Tilikete, et al., 2018a; Bonato et al., 2019; also see the **Supplementary Materials**). Procedures for testing fixed effects or for dealing with convergence problems were also unchanged with respect to our previous approaches (Blini, Desoche, et al., 2018b; Blini, Tilikete, et al., 2018a).

We tested the role of the following three factors (fixed effects) and their interactions: Congruency (2 levels: Congruent, Incongruent), Condition (3 levels: Potential Gain, PG; Neutral, N; Potential Loss, PL), and GVS (3 levels: sham; Right-Anodal, RA; Left-Anodal, LA). The random slope of Direction of the central arrow (and response effector, i.e. index or middle finger) was tested as well. We did not foresee, in the context of our study, any potential interest of this variable when tested as fixed effect. However, random slopes may account for, if selected in the models, part of the variability in the data. To summarize, the order in which we tested random slopes was the following: Congruency, Condition, GVS, and Direction.

The key test of this study was the three-way interaction Congruency by Condition by GVS. In addition, the following tests were part of our outcome-neutral quality checks (see below): Congruency, Condition, and Condition by Congruency. These four tests constitute our family of tests of interest (Cramer et al., 2016); the respective p-values were corrected for false discovery rate (Benjamini & Hochberg, 1995) separately from all remaining tests, i.e. exploratory tests. All p-values had to be significant at the 0.034 alpha threshold in light of sequential testing procedures. Any follow-up t-test comparison for significant effects was corrected for false discovery rate. Robustness checks and exploratory analyses were also envisaged following the results of this first analysis, and not pre-registered.

The SVV task was analyzed as in Blini et al. (2018a), and with the same statistical pipeline outlined above. First, extreme responses (exceeding ±2.5 standard deviations from the subject-specific mean) were discarded (0.01%). Then, we tested the fixed factors GVS (3 levels: sham; Right-Anodal, RA; Left-Anodal, LA) and Starting Orientation (2 levels: clockwise, counter-clockwise); this is also the order in which we tested random slopes. We predicted a large, yet not of interest, effect for Starting Orientation – lines initially displayed as tilted clockwise associated with a clockwise response bias. P-values for GVS and the twoway GVS by Starting Orientation interaction were adjusted for false discovery rate (Benjamini & Hochberg, 1995).

While the SVV task represents the elective control for the effectiveness of GVS, we also evaluated subjective sensations following each session (e.g. subjective vestibular effects, distress or motivation, as in Blini et al., 2018a). Each question was presented on a computer screen, above a horizontal line representing a visual scale continuum. Subjects graded their experience by means of mouse-clicks on the line. The standardized displacement from the objective center of the segment (i.e. −100% for the leftmost end, 0% for the exact center, +100% for the rightmost end) was used as the dependent variable for each question. The responses were submitted, for each of the 17 questions (original items in the **Supplementary Materials**), to a within-subject ANOVA with GVS as independent variable. We did not register in advance any specific prediction. However, three items will specifically assess unspecific (non-vestibular) distress: “I felt hitching”, “I felt burning”, and “I felt distress”. For these three items we required no significant effects in the aforementioned ANOVA (quality check 2, see below).

### 2.5 Predictions

We anticipated three possible scenarios for the three-way interaction Congruency by Condition by GVS, graphically depicted in **Figure 2C**. Our a priori hypothesis was that GVS, especially Right-Anodal, could decrease interference costs in PG conditions, in light of the decreased sensitivity to rewards (Blini, Tilikete, et al., 2018a); interference costs in the PL condition, instead, could be enhanced by either GVS montage with respect to sham (Scenario 3).

### 2.6 Outcome-neutral quality checks

Outcome-neutral quality checks are summarized in **Table 1**.

**Table 1:** Summary of outcome-neutral quality controls.

| Quality Control | Test | Invalidate results? | Outcome |
| --- | --- | --- | --- |
| The vestibular system is effectively perturbed | SVV task: the main effect of GVS or the GVS by Starting Side interaction is significant. At least one contrast indicates the following pattern: Left-Anodal < sham < Right-Anodal | √<br>No evidence for a successful vestibular stimulation | √<br>The subjective vertical was displaced according to the position of the anode: Left-Anodal < sham < Right-Anodal |
| Effects are not due to unspecific effects of | Questionnaires: no significant differences must be observed in items of the questionnaire | √<br>No evidence for vestibular | √<br>Distress or other |
| electric stimulation | evaluating unspecific sensations (e.g. distress, burning or itching sensation, pain...). Differences may be expected in items assessing vestibular-specific sensations (e.g. illusion of body movement) | specificity of the effects | unspecific sensations did not differ between GVS conditions and sham |
| The task evokes interference costs due to conflicting visual inputs | FT task: main effect of Congruency showing improved performance for congruent vs. incongruent trials | √<br>No evidence for response monitoring resources to be deployed | √<br>Responses to incongruent trials were slower and less accurate |
| Motivation is effectively manipulated | FT task: main effect of Condition (improved performance for PG and/or PL vs. Neutral) OR Condition by Congruency interaction (interference costs enhanced in PG and/or PL vs. Neutral) | X<br>Provided motivational assets are affected by GVS (main hypothesis), otherwise no evidence for an effective manipulation of motivational levels | √<br>Enhanced costs when value was at stake, especially for PL conditions; three-way interactions were involved |

#### Vestibular stimulation

The effectiveness of the vestibular stimulation had to be confirmed by one independent task (SVV). GVS was expected to tilt the SVV towards the site of anodal stimulation (Blini, Tilikete, et al., 2018a; Lenggenhager et al., 2007). We required at least one significant contrast along the Left-Anodal < sham < Right-Anodal pattern, as in Blini et al. (2018a). This was necessary in order to ascribe any behavioral effect to vestibular mismatches.

#### Vestibular specificity

In addition to the previous point, we needed to rule out the possibility that unspecific factors (e.g. distress, itching) were differing between the sham and GVS conditions. Participants’ responses to the questionnaire administered at the end of each session, thus, had to be comparable across GVS conditions (Blini, Tilikete, et al., 2018a).

#### Conflict and interference costs

The presence of interference costs – i.e. better performance for congruent vs incongruent trials – had to validate our modified version of the flankers task, and confirm that monitoring resources and parallel processing of irrelevant information were deployed.

#### Effects of motivation

Points at stake, signaled by the color of flanking arrows, had to modulate participants’ performance, as to ensure that different levels of motivation were indeed deployed in the task. This could happen either through improved performance in PG and/or PL conditions with respect to the neutral one (i.e. main effect of Condition), or – as could be predicted on the basis of studies assessing the modulatory effect of reward on interference costs (Krebs et al., 2010) – through enhanced interference costs for these conditions (e.g. hampered performance to incongruent trials). Given the aim of the present study, we did not consider fatal the lack of this quality check, but only in the presence of a meaningful modulation of motivational assets by GVS. For example, Scenario 1 in **Figure 2C** predicts large interference costs for the sham condition in face of smaller costs with concurrent GVS: in this case it would be conceivable that the two-way interaction Congruency by Condition (thus collapsed across GVS levels) may fail to reach our significance criterion, precisely because moderated by GVS; we would therefore observe only a three-way interaction, suggesting that motivational assets are indeed deployed, though modulated by GVS.

## 3. Results

### 3.1 Outcome neutral quality checks

#### Subjective Visual Vertical (SVV)

When assessing the SVV, we found, as predicted, a large effect of Starting Side *χ^2^*_(1)_= 66, *p_fdr_*< 0.001, *η_p_^2^*= 0.59), lines originally tilted in the clockwise direction being associated with a more pronounced clockwise bias. Additionally, the main effect of GVS was large and significant (*χ^2^*_(2)_= 62.76, *p_fdr_*< 0.001, *η_p_^2^*= 0.42). Post-hoc t-tests showed that all GVS conditions differed: Left-Anodal was associated with a bias in the counter-clockwise sense with respect to the sham (*β*= 0.166, SE= 0.066, z= 2.52, *p_fdr_*= 0.012); Right-Anodal, instead, was associated with a bias in the clockwise sense with respect to sham *φ*= 0.334, SE= 0.056, z= 6.01, *p_fdr_*< 0.001). GVS and Starting Side did not interact *χ^2^*_(2)_= 0.72, *p_fdr_*= 0.7). Results are depicted in **Figure 3A**.

#### Questionnaires

Across all 17 items in the questionnaire, we did not highlight GVS-specific differences for 16. In particular, there were no differences for questions assessing unspecific effects of an electric brain stimulation, e.g. “I felt itching”, “I felt burning”, or “I felt distress” (all *F*< 1.33, all *p*> 0.27). Likewise, nausea sensations did not differ from sham (*F*_(1.97, 80.58)_= 0.56, *p*= 0.57). However, when assessing participants’ illusions of their body moving along the sagittal axis (backward – forward) we found a significant effect of GVS (*F*_(1.75, 71.65)_= 6.74, *p*= 0.003, *η_p_^2^*= 0.14). Left-Anodal GVS was associated, with respect to both sham and Right-Anodal, to illusions of participants’ bodies displacing forward (all t > 2.54, all *p_fdr_*< 0.022). Degrees of freedom with decimal points denote Greenhouse-Geisser correction for violation of sphericity.

#### FT task

We found a main effect of Congruency for both accuracy (*χ^2^*_(1)_= 155.76, *p_fdr_*< 0.001, *η_p_^2^*= 0.81) and RTs (*χ^2^*_(1)_= 65.6, *p_fdr_*< 0.001, *η_p_^2^*== 0.64). Incongruent trials were less accurate and slower than congruent trials. The main effect of Condition was also significant for both accuracy (*χ^2^*_(2)_= 60.64, *p_fdr_*< 0.001, *η_p_^2^*= 0.4) and RTs (*χ^2^*_(2)_= 63.47, *p_fdr_*< 0.001, *η_p_^2^*= 0.32). However, post-hoc contrasts showed a dissociation between the two measures: for accuracy, both PG (9= 0.248, SE= 0.043, z= 5.74, *p_fdr_*< 0.001) and PL (*β*= 0.277, SE= 0.049, z= 5.63, *p_fdr_*< 0.001) conditions resulted in less accurate responses with respect to neutral trials; for response times, PG trials were associated to faster responses with respect to both neutral (*β*= 4.09, SE= 0.91, z= 4.5, *p_fdr_*< 0.001) and PL trials (*β*= 4.76, SE= 0.7, z= 6.8, *p_fdr_*< 0.001). Furthermore, while Congruency and condition did not interact when assessing accuracy (*χ^2^*_(2)_= 3.63, *p_fdr_*= 0.16), the two-way interaction was highlighted for RTs (*χ^2^*_(2)_= 10.09, *p_fdr_*= 0.009, *η_p_^2^*= 0.12). Specifically, interference costs (i.e. the difference between congruent and incongruent trials) were enhanced for PL trials with respect to both PG (*β*= 4.15, SE= 1.55, z= 2.67, *p_fdr_*= 0.011) and neutral trials (*β*= 4.72, SE= 1.57, z= 3, *p_fdr_*= 0.008); costs did not differ between PG and neutral trials (*β*= 0.57, SE= 1.37, z= 0.42, *p_fdr_*= 0.67). These tests are depicted in **Figure 3B** (accuracy) and **Figure 3C** (RTs).

**Figure 3:**
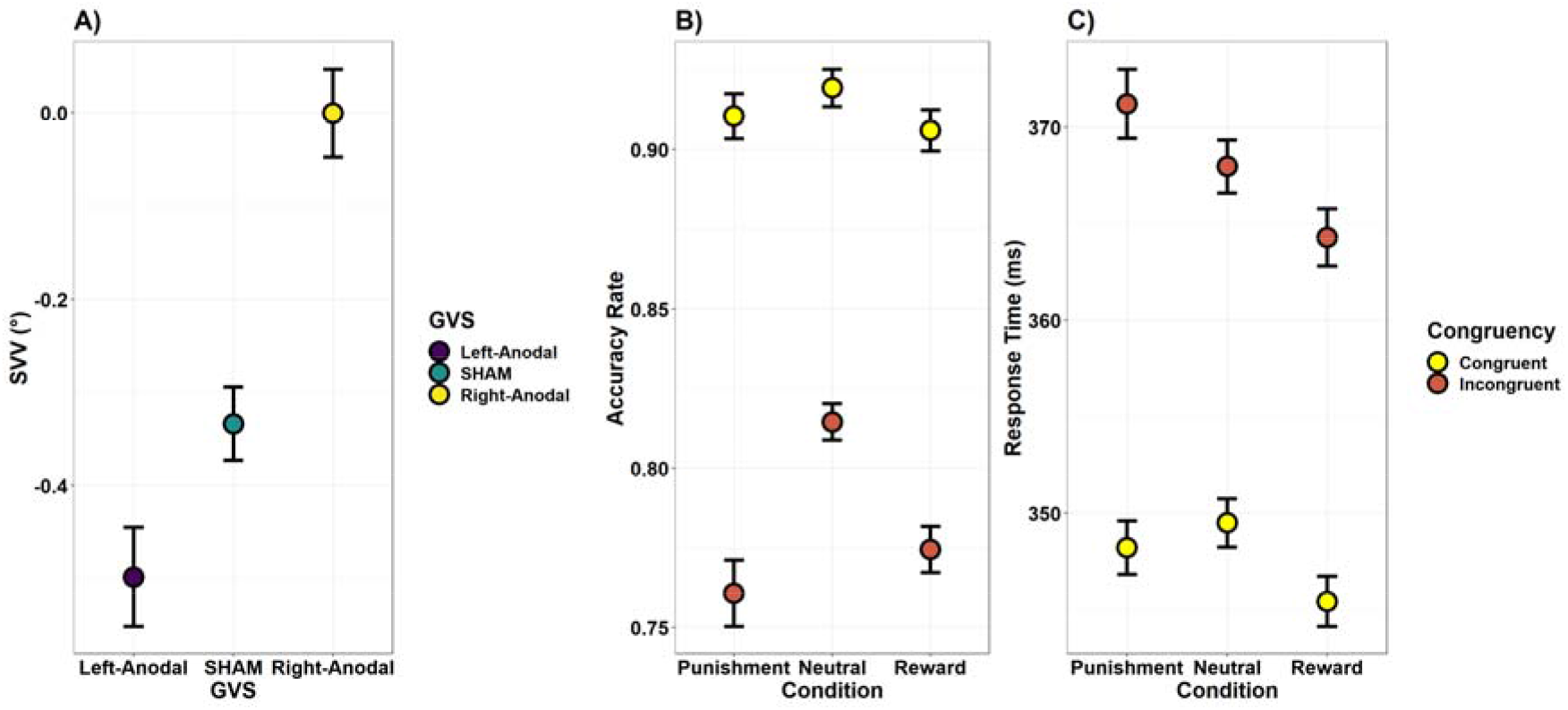
**A) Subjective Visual Vertical.** The subjective perception of verticality was effectively modulated by GVS. Left-Anodal GVS was associated with a counter-clockwise tilt of the SVV, clockwise for Right-Anodal, with respect to sham. **B) Condition and Congruency - Accuracy.** Interference costs (Congruent vs. Incongruent trials) were firmly highlighted. Both Potential Gain (Reward) and Potential Loss (Punishment) conditions, interestingly, were associated with decreased accuracy. Thus, in the FT task, motivational cues, attached to distractors, were actually impairing performance. **C) Condition and Congruency - Response Times.** Interference costs (Congruent vs. Incongruent trials) were firmly highlighted. Differently from accuracy, Potential Gain (Reward) conditions were overall faster than Neutral and Potential Loss trials. However, only Potential Loss trials were interacting with congruency such that interference costs were increased in this condition with respect to both Neutral and Potential Gain trials. Thus, Reward and Punishment conditions seemed to dissociate for response times. The fact that Punishments led to slower responses than or equal to Neutral trials seems to point against the possibility that participants were slowing down selectively and strategically when no reward was at stake. Error bars depict within-subjects standard errors of the mean (Morey, 2008).

#### Discussion

All the outcome-neutral quality checks were fulfilled. First, the SVV task provided independent evidence that our GVS stimulation was, indeed, altering vestibular processing. The subjective visual vertical was tilted toward the site of anodal GVS stimulation, possibly in light of a modulation of the input originating from the otoliths. Participants also reported, after Left-Anodal GVS, having perceived their body moving forward in space. Albeit this is compatible with a vestibular stimulation, the direction of this illusion is unusual for a bilateral montage. Displacements along the anterior-posterior axis may be expected for antero-posterior montages (Aedo-Jury, Cottereau, Celebrini, & Severac Cauquil, 2019), rather than for lateral ones. This illusion may have been genuinely prompted by the (mixed) vestibular signals that participants experienced, or could simply reflect difficulties in accurately discriminating or recall (after the stimulation) the sensations experienced. That said, all other items indicated that GVS was not different from sham when assessing potential confounds that may follow electric brain stimulations, such as overall discomfort or painful sensations. This is important because it allowed us to circumscribe the experimental effects more precisely to a vestibular perturbation. Finally, we have found, in the FT task, all the requisites for us to assess a meaningful modulation by GVS: conflict costs, in terms of impaired responses in incongruent trials (Congruency effect), and motivational effects. The latter ones, interestingly, caused an impaired performance when value was at stake (less accurate responses). This is coherent with the view that distractors may gain in salience when value is attached to them, and hamper the performance to the relevant task. However, PL and PG conditions seem to dissociate when assessing response times: PG trials were overall faster, suggesting a motivational performance boost; PL trials, on the other hand, were not faster than neutral trials, but they interacted instead with Congruency such that the observed conflict costs were enhanced (i.e. responses were faster in congruent trials but slower in incongruent trials, **Figure 3C**). This accurately reflects the pattern depicted in Figure 1C as an increase, with respect to neutral trials, of interference costs in PL (punishment) conditions; with respect to our a priori predictions, however, we have found no evidence of such increase in PG (rewarded) conditions.

### 3.2 The vestibular system, conflict, and motivation

This section reports the results of the main tests of interest, namely the three-way interaction Congruency by Condition by GVS for both accuracy (**Figure 4A**) and RTs (**Figure 4B**). Descriptive statistics are reported in **Table 2**.

**Figure 4:**
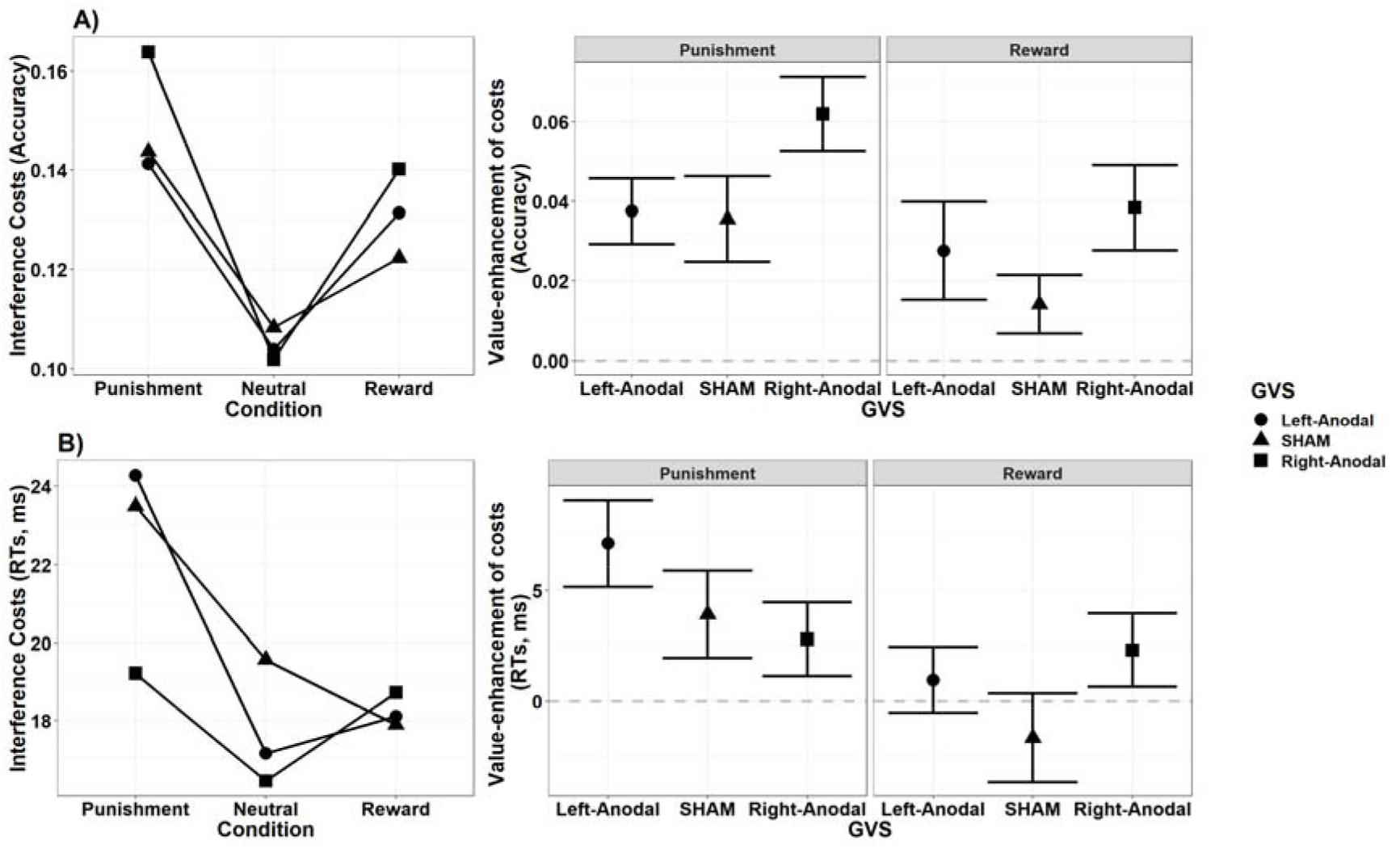
Results of the main tests of interest, namely the three-way interaction Congruency by Condition by GVS for both accuracy (**A)**) and RTs (**B)**). The leftmost plots of each panel show interference costs for each GVS and Reward condition, depicted as in **Figure 1C**. In the rightmost plots of each panel, instead, we use interference costs in Neutral trials (dashed gray line) as a baseline to assess how different Reward conditions and GVS can modulate them. We have found weak evidence that: Right-Anodal GVS increases, when assessing accuracy, the interference costs associated with PL trials; Left-Anodal GVS increases, when assessing RTs, the interference costs associated with PL trials (with respect to Right-Anodal GVS). Results should, however, be taken with caution because: effects sizes were very small (about half the expected size); a range of exploratory robustness checks (presented in section 3.3) failed to fully corroborate the results from this main, pre-registered analysis. Error bars depict within-subjects standard errors of the mean (Morey, 2008).

**Table 2:** Descriptive statistics.

| **GVS** | **Condition** | **Congruency** | **Accuracy (%)** | **SD** | **Response Times (ms)** | **SD** |
| --- | --- | --- | --- | --- | --- | --- |
| Left-Anodal | PL | Congruent | 90.65 | 8.64 | 345.7 | 34.01 |
|  |  | Incongruent | 76.52 | 11.96 | 369.98 | 51.76 |
|  | Neutral | Congruent | 91.71 | 8.43 | 347.63 | 36.18 |
|  |  | Incongruent | 81.33 | 10.56 | 364.77 | 48.3 |
|  | PG | Congruent | 91.01 | 10.01 | 343.75 | 35.67 |
|  |  | Incongruent | 77.87 | 10.94 | 361.86 | 50.73 |
| sham | PL | Congruent | 90.37 | 9.88 | 347.07 | 34.29 |
|  |  | Incongruent | 75.99 | 13.26 | 370.58 | 52.12 |
|  | Neutral | Congruent | 91.87 | 9.07 | 348.52 | 35.81 |
|  |  | Incongruent | 81.04 | 11.62 | 368.09 | 52.26 |
|  | PG | Congruent | 89.77 | 10.45 | 344.55 | 35.7 |
|  |  | Incongruent | 77.54 | 11.9 | 362.44 | 50.88 |
| Right-Anodal | PL | Congruent | 92.11 | 7.83 | 349.28 | 29.91 |
|  |  | Incongruent | 75.73 | 11.8 | 368.5 | 43.62 |
|  | Neutral | Congruent | 92.2 | 7.45 | 350.11 | 30.63 |
|  |  | Incongruent | 82.01 | 11.2 | 366.52 | 40.57 |
|  | PG | Congruent | 91.01 | 8.36 | 345.04 | 30.35 |
|  |  | Incongruent | 76.98 | 12.01 | 363.78 | 42.07 |

When assessing accuracy, the interaction was not significant according to our pre-defined criteria (*χ^2^*_(4)_= 9.62, *p_fdr_*= 0.063, *η_p_^2^*= 0.05), though it was showing a trend at the uncorrected level (p= 0.047). We decided to perform follow-up tests regardless, to better frame the source of this tendency, and noticed that PL trials were associated with larger interference costs with respect to Neutral trials only in the Right-Anodal GVS condition (*β*= 0.386, SE= 0.125, z= 3.09, *p_fdr_*= 0.006). We then moved to assessing RTs. Here, the interaction was significant (*χ^2^*_(4)_= 10.75, *p_fdr_*= 0.029, *η_p_^2^*= 0.04), albeit the associated effect size was small. Follow-up t-tests pointed toward the fact that, in PL conditions, interference costs were enhanced comparatively more for the Left-Anodal condition with respect to the Right-Anodal one (*β*= 6.7, SE= 2.88, z= 2.33, *p_fdr_*= 0.06 – uncorrected *p*= 0.02), though not when compared to sham (*β*= 1.5, SE= 3.06, z= 0.49, *p_fdr_*= 0.62).

Besides the tests reported under “Outcome-neutral quality checks”, no other effect or interaction was proven significant when assessing either accuracy or RTs (all *p_fdr_* > 0.14).

### 3.3 Robustness checks

This part was not pre-registered. The aim of the analyses in this section was to probe the main results of our study, i.e. concerning the three-way interaction Congruency by Condition by GVS, through different statistical techniques in order to verify their robustness. For a complete overview, the reader is referred to the **Supplementary Materials**.

We performed 2 (Congruency: congruent, incongruent) x 3 (Condition: PL, PG, N) x 3 (GVS: Right-Anodal, Left-Anodal, sham) repeated measures ANOVAs on the data from the FT task, preprocessed as in the main analyses. Degrees of freedom with decimal points indicate that Greenhouse-Geisser correction for violation of sphericity was applied. Accuracy data were previously arcsine square root transformed in order to use proportions as dependent variable (but note that, with respect to logistic link-functions, this approach decreases the power of a test, Warton & Hui, 2011). The three-way interaction Congruency by Condition by GVS was not significant when assessing accuracy (*F*_(3.37, 138.3)_= 2.1, *p*= 0.1) or RTs (*F*_(3.53, 144.92)_= 1.91, *p*= 0.12). The latter analysis did not change significantly when target Direction was added among the factors (p= 0.09) or when different thresholds for the removal of outliers were adopted (minimum p= 0.09 when 2.6 standard deviations was used as criterion). Because effects were subtle, but to some extent similar between accuracy and RTs, we computed a simple Efficiency index (Accuracy Rate divided by RTs, in seconds). Larger values indicate better performance. However, in light of the high variability of these values, the three-way interaction (Congruency x Condition x GVS) was not significant either (*F*_(3.24, 132.64)_= 1.52, *p*= 0.21). We therefore proceeded with normalizing the data to a more stable baseline: for each subject, we first computed the interference costs (the difference between congruent and incongruent trials) for each GVS and Condition, and then referenced these costs to the Neutral condition (situation depicted in the rightmost plots of **Figure 4**). Thus, larger values in this context indicate larger enhancement of interference costs when value was at stake, with respect to a Neutral baseline. We probed the same post-hoc contrasts which resulted significant in the preregistered approach, which was based on mixed models, through paired t-tests on these values. When assessing accuracy, we have found that Right-Anodal GVS yielded, in PL trials, larger costs than both sham (*t*_(41)_= 2.32, *p*= 0.026) and Left-Anodal GVS (*t*_(41)_= 2.09, *p*= 0.043), though only at the uncorrected significance level. When assessing RTs, instead, the difference between Right-Anodal and Left-Anodal GVS when potential losses were at stake was not significant (*t*_(41)_= 1.73, *p*= 0.091).

### 3.4 Exploratory analyses

#### Drift Diffusion Models

Another way to link accuracy rate and response time via one theory-grounded framework is through Drift Diffusion Models (**DDM**, Ratcliff, 1978). DDM assume that information is sampled continuously from the environment and, in the case of a two forced-choices task, one response is produced when a critical threshold is reached. Thus, at least two parameters are of interest: the drift rate, indexing the direction and speed of sensory accumulation, and the boundary separation parameter, indexing the amount of information that is necessary in order to produce a response. The first one has been classically associated with task difficulty – e.g. the amount of cognitive load, which hampers the speed by which information is gathered – whereas the second is thought to refer to speed-accuracy tradeoffs and impulsivity – larger boundary separation suggesting a more conservative threshold and response style. DDM approaches for the Flankers task have been discussed before (Fischer et al., 2018; White et al., 2011). We fitted one such DDM via the fast-dm software (Voss & Voss, 2007), choosing the Kolmogorov-Smirnov minimization procedure and setting the variance parameters (drift rate, starting position) to 0 (Voss, Voss, & Lerche, 2015). We used accuracy coding such that the upper response boundary represented a correct response. The procedure recovered, for each subject, the drift rate and boundary separation parameters, which were submitted to 2 (Congruency: congruent, incongruent) x 3 (Condition: PL, PG, N) x 3 (GVS: Right-Anodal, Left-Anodal, sham) repeated measures ANOVAs. For the drift rate parameter, we observed a main effect of Congruency (*F*_(1,41)_= 195.64, *p*< 0.001, *η_p_^2^*= 0.83) and Condition (*F*_(1.98,81.36)_= 23.63, *p*< 0.001, *η_p_^2^*= 0.37), as well as the interaction between the two (*F*_(1.79,73.29)_= 4.26,*p*=0.02, *η_p_^2^*= 0.09). Incongruent trials were associated with slower drift rates than congruent ones (*t_(41)_*= 13.99, *p_fdr_*< 0.001). Interference costs were particularly pronounced, in addition, when value was at stake, thus for PL (*t_(41)_*= 6.12, *p_fdr_*< 0.001) and PG trials (*t_(41)_*= 6.05, *p_fdr_*< 0.001) with respect to Neutral, without differences between PL and PG (*t_(41)_*= 0.03, *p_fdr_*= 0.98). This is coherent with the view that both distractors and value information have detrimental effects on the task at play, hampering the gathering of relevant information. However, Condition and Congruency interacted, and PL trials only were associated with increased interference costs (*t_(41)_*= 2.74, *p_fdr_*= 0.027) with respect to Neutral trials. Thus, this finding seems to suggest that PL trials may be more effective in capturing attention and causing distraction with respect to PG ones, corroborating the PL-associated interference cost unveiled with the pre-registered LMEM analysis. There was a weak tendency for this effect to be larger with concurrent Right-Anodal GVS (PL vs Neutral, Left-Anodal vs. Right-Anodal: *t_(41)_*= 2.11, *p*= 0.04; PL vs Neutral, sham vs. Right-Anodal: *t_(41)_*= 2.06, *p*= 0.046); Right-Anodal was, indeed, the only GVS condition in which Congruency and Condition interacted (*F*_(1.95,79.97)_= 6.06, *p*= 0.004, *ηp^2^*= 0.13), but not Left-Anodal (*F*(1.85,75.86)= 0.88, *p*= 0.41) or sham (*F*_(1.96,80.44)_= 1.43, *p*= 0.24). However, the corresponding three-way interaction failed to reach significance (*F*(3.43,140.77)= 1.86, *p_fdr_*= 0.13). Results are depicted in **Figure 5**. When assessing the boundary separation parameter, interestingly, we only found a main effect of Condition (*F*_(1.88,76.92)_= 28.86, *p*< 0.001, *η_p_^2^*= 0.41): PG trials were associated with lower values than PL trials (*t_(41)_*= 4.99, *p_fdr_*< 0.001), which were in turn associated with lower values than Neutral trials (*t_(41)_*= 2.72, *p_fdr_*= 0.01). Thus, responses were more impulsive when value was at stake, potential gain trials being associated with the most liberal response bias.

**Figure 5:**
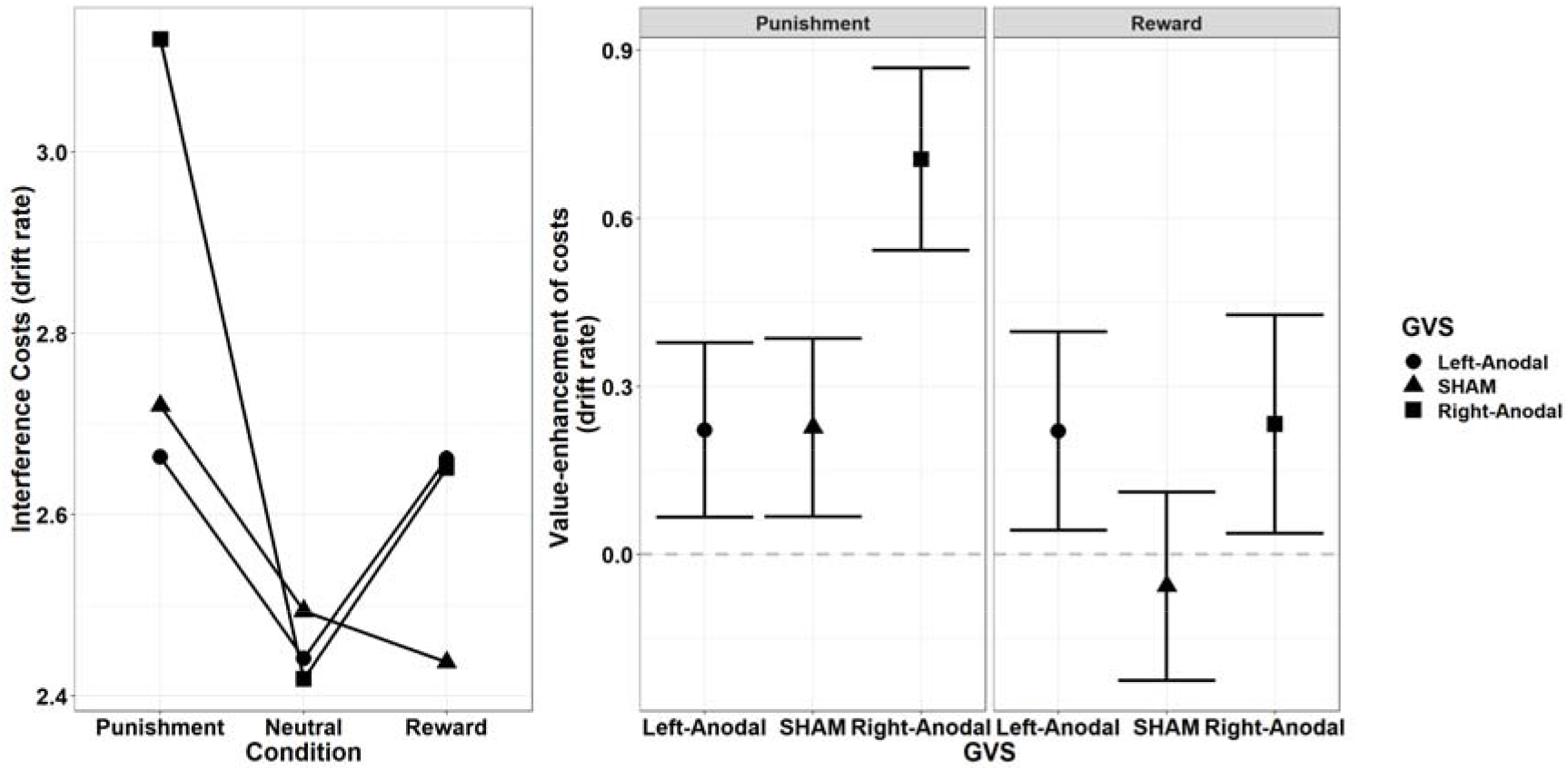
We fitted a Drift Diffusion Model (DDM) to the data, and recovered the drift rate parameter, indexing the speed at which sensory information is gathered from the environment. Such speed was slower for incongruent trials and when value was at stake, showing that irrelevant information was indeed causing distraction and hampering the performance to the relevant task. These interference costs were larger for PL trials, suggesting that potential losses were more effective in capturing attention. We have additionally found some evidence for this effect to be more pronounced with Right-Anodal GVS. The leftmost plot shows interference costs for each GVS and Reward condition, depicted as in **Figure 1C**. In the rightmost plot, instead, we use interference costs in Neutral trials (dashed gray line) as a baseline to assess how different Reward conditions and GVS can modulate them. Error bars depict within-subjects standard errors of the mean (Morey, 2008).

### 3.5 Summary of results

All the quality controls planned for the study were fulfilled, enabling us to interpret the results from the FT task across GVS conditions. One first result was that PL and PG trials dissociated. While both had detrimental effects on behavioral performances, as assessed by accuracy, only PL trials led to enhanced interference costs when assessing RTs. Thus, value can have detrimental effects on performance, when it is attached to task-irrelevant information, possibly as result of increased attentional salience of the distractors. Potential losses, in this respect, seem to be more effective than potential gains, suggesting some prioritization. The main objective of this study, however, was to assess whether this interaction was further modulated by GVS. We have found some evidence in this regard:Right-Anodal marginally increased interference costs for PL trials when assessing accuracy, whereas Left-Anodal GVS was seemingly more effective when assessing response times. It must be said, however, that several post-hoc robustness checks could not corroborate the results of the main analyses, the p-values being close (~0.1) to significance but not comparable to LMEM results. Importantly, the effect size for the three-way interaction was rather small (*η_p_^2^*= ~0.05), half of what hypothesized for the a priori power analysis. Post-hoc power for our design equals to only 50% for such small effects (given our set of p-value corrections).

LMEM are, indeed, much more powerful than traditional approaches in their using all the available data (i.e. the single trial level). However, many researchers’ degrees of freedom are associated with their use. For example, one random effects matrix must be specified, and this choice heavily affects the results. Deficitary matrices are associated with extremely liberal LMEM results (high type 1 error rates); for this reason, a few authors suggested to simply exploit the most complex matrix of random effects available (justified by the experimental design, Barr, Levy, Scheepers, & Tily, 2013). However, this comes with a cost, and models that are over-parametrized tend not to converge and severely hamper power (type 2 errors, Matuschek, Kliegl, Vasishth, Baayen, & Bates, 2017). Indeed, we tried to fit the maximal models in our experiment for both accuracy and response times, but the models could not converge. Here, we have tried to reach a parsimonious solution through an objective, preidentified, pipeline. We are unable to state whether our results are due to a genuinely more powerful approach or to a suboptimal matrix of random effects. There are, however, also encouraging points: first, results are rather coherent for both accuracy and RTs, in their showing a pattern of increased value-specific interference during GVS; this is reaffirmed by exploratory analyses exploiting drift diffusion models, which organically link accuracy and RTs, showing indeed some role of Right-Anodal GVS in promoting attention to punishments. While this pattern roughly matches our predictions of an increased saliency – and therefore interference costs – for PL trials during visuo-vestibular mismatches, we found no evidence for a reduction of such costs for PG trials, which was also expected.

## 4. Discussion

### 4.1 Motivation, valence and conflict

Motivation – experimentally provided, for example, in the form of monetary rewards – can act by invigorating one’s action or willingness to exert an effort (Chong, Bonnelle, & Husain, 2016; Chong et al., 2015; Muhammed et al., 2016), or by sharpening cognitive processes such as attention and memory (Abe et al., 2011; Anderson et al., 2011; Chelazzi et al., 2013; Della Libera & Chelazzi, 2006, 2009; Engelmann & Pessoa, 2007; Hickey et al., 2010). However, recent developments also highlight instances in which rewards may hamper performance (Krebs et al., 2010; Marini, Berg, & Woldorff, 2015; Theeuwes & Belopolsky, 2012; Watson, Pearson, Theeuwes, Most, & Le Pelley, 2020), for example when attention is unduly captured by distracters that are related to rewards themselves, or in presence of conflicting information. In the present study, we report data showing both facets of motivation. The invigorating component was exemplified by overall faster response times when potential gains were at stake. The detrimental component, on the other hand, had a larger share, and we observed both decreased accuracy when gains and losses were at stake and a peculiar increase in interference costs specifically for potential losses.

While the bulk of brain circuits coding for rewards and punishments appears to be highly overlapping, signals associated with gains or losses are also differentially represented in the brain (Camara et al., 2009; Liu et al., 2011; Yacubian et al., 2006). Specialised brain areas originate a signal coding for relative reward magnitudes, which is thought to be exploited for behavioural optimization (Bush et al., 2002; Hernandez Lallement et al., 2014; Hickey et al., 2010; Sallet et al., 2007; van Steenbergen et al., 2012; Williams et al., 2004). A subset of these areas has also been associated with processes involving response monitoring, especially under conflict (Botvinick et al., 2004; Carter et al., 1998; Floden et al., 2010). Recent proposals suggest that the dorsal Anterior Cingulate Cortex (**dACC**), in particular, may code conflict as an aversive signal biasing behaviour away from irrelevant information and suboptimal outcomes (Botvinick, 2007; Inzlicht et al., 2015). One recent study, for example, found that the dACC includes both conflict- and value-related representations (Vermeylen et al., 2019): a multivariate algorithm trained to classify affective value (negative vs positive stimuli) could later classify above chance conflict (incongruent vs congruent trials) in an independent task within dACC and the pre-supplementary motor area. That only losses result in enhanced interference costs, thus, may indicate that the reported overlap is not merely evaluative in its functions. Rather, potential losses may consolidate and magnify neural signals related to conflict, and thus the associated behavioral signatures. There are at least two possible mechanisms that may frame this possibility. First, potential losses may be more effective than potential gains in capturing attention: the processing of task-irrelevant features (the flankers’ direction) would also be enhanced as well as a consequence, causing a more pronounced interference. Second, one additional source of conflict may arise between an intrinsic tendency to associate losses with avoidance and the need to provide a response as fast as possible (Carsten et al., 2019); gains, evoking approach tendencies instead, would not be associated with such additional mismatch. We tend, however, to favor the first option as it best accommodates our finding that incongruent trials were, indeed, slower, whereas congruent ones were slightly faster than the respective neutral condition. An unspecific source of conflict and avoidance prompted by losses should, instead, equally affect congruent and incongruent trials, causing response times to be overall slower. Drift diffusion models have shown, indeed, that responses were generally less impulsive when potential losses were at stake with respect to potential gains, which is compatible with this view. However, potential losses were associated, on the contrary, to more impulsive responses with respect to neutral trials, and thus impulsivity alone is unlikely to explain the increased interference costs only observed for losses. Marginally faster response times for congruent trials, instead, are compatible with the attentional account because the enhanced processing of the distractors’ direction would, in this specific condition, hamper comparatively less the response to the primary task. In other words, part of the sensory information that is gathered from the distractors could, in a congruent condition, be more easily “recycled” and transferred to the task-relevant response. Indeed, this interaction between losses and congruency is best captured by the drift rate parameter, indexing the speed at which sensory information is gathered from the environment.

The notion that potential losses may be more effective than potential gains in driving decision-making is not new (see the prospect theory: Kahneman & Tversky, 1979), though experimental evidence is to date mixed. Several studies found similar effects on motivation for reward and punishment (Nissens et al., 2017; Wentura et al., 2014) or even seemingly more pronounced effects for the former (Carsten et al., 2019); recent research, on the other hand, describes instances in which loss aversion can dominate over reward seeking (Liao et al., 2020; Massar et al., 2020). Liao and colleagues (2020), in particular, specifically assessed the influence of threat and aversive motivation (i.e. the delivery of electric shocks) on conflict processing in the context of a Stroop task (Krebs et al., 2010). The authors reported that, in contrast with potential gains, aversive motivation was associated with impaired performance in the task which, like in the present study, was counterproductive for participants as it resulted in increased occurrence of electric shocks; in addition, interference costs were increased for incongruent trials, though in a manner qualitatively similar to that observed in potential gain blocks (Liao et al., 2020).

All in all, thus, our results seem to corroborate the notion that potential losses may drive decision-making more powerfully than potential gains, the inconsistencies found in literature being explained by a different sensitivity of the behavioral tasks or by methodological constraints (e.g., intermixed presentation of PG and PL trials versus in separate blocks).

### 4.2 The vestibular system, value, and conflict

The role of the vestibular system in modulating motivational and monitoring resources remains, at state, elusive. The present study could only provide weak and ambiguous evidence for GVS to further qualify the interaction between value and conflict. If we were to rigidly apply our pre-registered criteria in evaluating this three-way interaction, we should conclude that Left-Anodal GVS may increase interference costs for potential losses (when assessing response times). It would be therefore tempting to discuss results in terms of hemispheric asymmetries for the processing of valence. Left hemisphere structures have been proposed to be dominant for approach behavior/positive emotions, and right hemisphere structures to be dominant for avoidance behavior/negative emotions (Berkman & Lieberman, 2009; Canli et al., 1998). It seems therefore fitting that Right-Anodal GVS, which activates comparatively more left hemisphere structures (Lopez et al., 2012; zu Eulenburg et al., 2012), may specialize for rewards (Blini, Tilikete, et al., 2018a) whereas Left-Anodal GVS, which activates mainly right hemisphere structures, may preferentially affect punishments. Though intriguing, however, any conclusion would be premature as a number of post-hoc analyses failed to support this finding. In addition, in our previous study, using a spatial cueing paradigm (Blini et al., 2018a), Left-Anodal GVS showed an effect in the same direction as that of Right-Anodal GVS, and thus decreased sensitivity to rewards as well, though to a lesser extent. Finally, in the present study we have also found hints suggesting that Right-Anodal GVS may enhance interference costs for potential losses. While robustness checks were, also in this case, far from providing a clear cut answer, this latter result was seemingly more robust internally. First, LMEM and robustness checks were roughly comparable, though the latter yielded post hoc contrasts that were only significant at the uncorrected level. Second, drift diffusion models, and the drift rate parameter specifically, could highlight a specific effect for Right-Anodal, but not Left-Anodal GVS, in the processing of losses. This pattern roughly matched our predictions (depicted in **Scenario 3, Figure 2C**) of an increased saliency – and therefore interference costs – for PL trials during visuo-vestibular mismatches, although the effect was much smaller than what foreseen; on the other hand, we have found no evidence for a reduction of such costs for PG trials, which was also expected across all a priori scenarios.

It should be noted that, unlike losses, gains did not yield an increase of interference costs; one possibility, thus, is that this supposed interaction is not something that could have been meaningfully modulated in first place. Rather, one may wonder why GVS did not modulate the response time advantage for gains, which we discussed above as reflecting an increase in response vigor due to the promise of rewards. One first possibility is that our previous study may simply report a false positive finding. The study was pre-registered as well: this process seems to work as intended in increasing the likelihood for null findings to be published (Allen & Mehler, 2019), and therefore mitigate publication bias, but it is not by itself a miracle cure for false positives, which are intrinsic to inferential statistics based on error rate probabilities. That said, the response times advantage reported in the present study is much more subtle than the one we previously described, obtained through a different task (a spatial cueing paradigm, Blini et al., 2018a). Furthermore, the Flankers task adopted in this study was explicitly meant to probe an instance in which value actually impairs behavioral performance; indeed, in the FT, gains were also *decreasing* performances in terms of discrimination accuracy. One could therefore simply argue that the effect of potential gains was, in the current context, too weak and ambiguous to be effectively modulated.

### 4.3 Limitations of the present study

In the present study, value information was attached to distractors, which were presented on screen simultaneously to targets. Participants’ responses occurred, on average, around 340-370 ms post-stimulus. This is a very short time window, which probably only allowed us to frame effects that are more likely to be related to very low-level, perceptual differences induced by the processing of value. Having observed such differences, for example in the processing of gains vs losses, in such a limiting setting is remarkable, and indicates that value can affect (and hamper) human performance very quickly, and starting with very basic sensory processes. Indeed, our findings were best captured by a difference in the speed of accumulation of evidence (the drift rate parameter), which is thought to reflect precisely sensory processing of the environment. On the other hand, we may have missed effects originating from late processes or components. In our previous study (Blini, Tilikete, et al., 2018a), the spatial cue signaling reward preceded the target of about 700 ms. More high-level (e.g. strategical) effects of reinforcers, in addition to perceptual ones, may be unveiled by allowing a more thorough processing of value. This, in turn, may increase the likelihood to induce and observe modulations of value-specific effects, for example by GVS.

Another aspect worth of consideration is the choice of the time threshold imposed to participants. One such threshold is important as it helps to highlight effects of value that are only present when the outcome of a trial is not certain, and the reinforcers are more likely to be obtained if comparatively more effort is deployed for the response. Differently from our previous approach (Blini et al., 2018a), this threshold was not fixed (e.g. to 500 ms), but adaptive, and it was meant to provide similar levels of task difficulty across all subjects. Participants were continuously pushed by the algorithm to produce responses faster than 75% of their overall responses. This context was overall stricter, and, in hindsight, may have caused participants to reach the limits of their performances (i.e. floor effect) sooner during each session. In addition, this could have introduced some powerful extrinsic (though orthogonal to all GVS and Conditions) source of motivation, of such a large magnitude to partly conceal that induced by reinforcers.

Another remark worth mentioning is that we used, for the present task, fixed color-value associations. Considering that the task did not involve statistical learning (i.e., color-value associations were explicit), we sought to minimize any potential conflict arising at the semantic level, as to avoid conditions of uneven strength of the associations. As the mesolimbic reward system may be differentially tuned to different color wavelength (Hu et al., 2020), however, this may have introduced perceptual biases effectively defeating the purpose.

Last but not least, here we choose to present potential loss and potential gain trials in an intermixed, balanced fashion. Compared to their administration in separate blocks (e.g., Liao et al., 2020), this may have prompted a competition, more or less explicit, between the two valences, resulting in one being favored and prioritized over the other, perhaps depending on individual biases and personality traits. Measuring and accounting for such individual susceptibilities may help clarify this issue in future studies.

### 4.4 Conclusions

In this paper, we report an asymmetry in the processing of gains and losses, with the latter seemingly more effective in capturing human attention. When negative reinforcers are attached to distractors, these stimuli appear to be processed more thoroughly, and the interference costs arising in presence of conflicting information are consequently enhanced. We also report some evidence for Right-Anodal galvanic vestibular stimulation to further increase the salience of losses, as suggested by even larger interference costs in this condition. This effect appears, within the present study, very weak, and thus of dubious relevance, although the behavioral task has shown much room for future improvements. While this pattern of results mimics the one we originally predicted as Scenario 3, we can only draw uncertain conclusions from our data, let aside any neural counterpart. However, these ambiguous results do stimulate new questions to be tackled in future endeavors: are vestibular links with motivation solid and practically relevant enough to be translated to addiction disorders or other clinical settings? Are the effects specific to valence or rather mediated by some other common feature? And what are the most likely neural counterparts of such effects: is there an asymmetry between GVS montages and the treatment of either punishments or rewards? Are predictive coding accounts, and the notion of prediction error signal specifically, useful to frame the interaction between the vestibular system and motivation? We call for high-powered, pre-registered studies in answering these questions, as a context in which null or ambiguous results are not stigmatized is paramount in reaching a real informed judgement.

## Author Contributions

**Elvio Blini:** Conceptualization; Data curation; Formal analysis; Funding acquisition; Investigation; Methodology; Project administration; Resources; Software; Validation; Visualization; Original draft; Review & editing.

**Caroline Tilikete:** Conceptualization; Methodology; Project administration; Supervision; Validation; Review & editing.

**Leonardo Chelazzi:** Conceptualization; Methodology; Supervision; Validation; Review & editing.

**Alessandro Farné:** Conceptualization; Methodology; Project administration; Supervision; Validation; Review & editing.

**Fadila Hadj-Bouziane:** Conceptualization; Methodology; Project administration; Supervision; Validation; Review & editing.

## Acknowledgments

EB was supported by: the European Union’s Horizon 2020 research and innovation programme (Marie Curie Actions) under grant agreement MSCA-IF-2016-746154; a grant from MIUR (Departments of Excellence DM 11/05/2017 n. 262) to the Department of General Psychology, University of Padova. AF is supported by the James S. McDonnell Scholar award. FHB received funding from the French National Research Agency (ANR) (ANR-14-CE13-0005-1). The study was performed within the framework of the LABEX CORTEX (ANR-11-LABX-0042) of the University of Lyon within the program “Investissements d’Avenir” (ANR-11-IDEX-0007) operated by the ANR. Funders have no role in study design, data collection and analysis, decision to publish, or preparation of the manuscript.

## Notes

https://osf.io/b9ezq/

### Competing Interest Statement

The authors have declared no competing interest.

### Summary of Updates

The discussion is now more detailed, with enhanced coverage of the literature. Limitations of the study are now flagged much more visibly. This was to accomodate reviewers' comments following peer-review at Cortex.

